# A novel bat coronavirus reveals natural insertions at the S1/S2 cleavage site of the Spike protein and a possible recombinant origin of HCoV-19

**DOI:** 10.1101/2020.03.02.974139

**Authors:** Hong Zhou, Xing Chen, Tao Hu, Juan Li, Hao Song, Yanran Liu, Peihan Wang, Di Liu, Jing Yang, Edward C. Holmes, Alice C. Hughes, Yuhai Bi, Weifeng Shi

## Abstract

The unprecedented epidemic of pneumonia caused by a novel coronavirus, HCoV-19, in China and beyond has caused public health concern at a global scale. Although bats are regarded as the most likely natural hosts for HCoV-19^1,2^, the origins of the virus remain unclear. Here, we report a novel bat-derived coronavirus, denoted RmYN02, identified from a metagenomics analysis of samples from 227 bats collected from Yunnan Province in China between May and October, 2019. RmYN02 shared 93.3% nucleotide identity with HCoV-19 at the scale of the complete virus genome and 97.2% identity in the 1ab gene in which it was the closest relative of HCoV-19. In contrast, RmYN02 showed low sequence identity (61.3%) to HCoV-19 in the receptor binding domain (RBD) and might not bind to angiotensin-converting enzyme 2 (ACE2). Critically, however, and in a similar manner to HCoV-19, RmYN02 was characterized by the insertion of multiple amino acids at the junction site of the S1 and S2 subunits of the Spike (S) protein. This provides strong evidence that such insertion events can occur in nature. Together, these data suggest that HCoV-19 originated from multiple naturally occurring recombination events among those viruses present in bats and other wildlife species.

## Text

Coronaviruses (CoVs) are common viral respiratory pathogens that primarily cause symptoms in the upper respiratory and gastrointestinal tracts. In 1960s, two CoVs, 229E and OC43, were identified in clinical samples from patients experiencing the common cold^3^. More recently, four additional human CoVs have been successively identified: severe acute respiratory syndrome coronavirus (SARS-CoV) in 2002, NL63 in late 2004, HKU1 in January 2005, and Middle East respiratory syndrome coronavirus (MERS-CoV) in 2012. However, only two betacoronaviruses (beta-CoVs), SARS-CoV and MERS-CoV, are able to cause severe and fatal infections, leading to 774 and 858 deaths, respectively, suggesting that beta-CoVs may be of particular concern to human health. In December 2019, viral pneumonia caused by an unidentified microbial agent was reported, which was soon identified to be a novel coronavirus^4^, now termed SARS-CoV-2 by the International Committee for the Taxonomy of Viruses^5^ and HCoV-19 by a group of Chinese scientists^6^. The number of patients infected with HCoV-19 has increased sharply since January 21, 2020, and as of March 2rd, 2020, more than 80,000 confirmed HCoV-19 cases have been reported, with >11,000 severe cases and >2900 deaths in China. By the end of January confirmed HCoV-19 cases were present in all the Chinese provinces and municipalities and at the time of writing the virus has been detected in over 60 countries.

An epidemiological survey of several HCoV-19 cases at an early stage of the outbreak revealed that most had visited the Huanan seafood market in Wuhan city prior to illness, where various wild animals were on sale before it was closed on January 1, 2020 due to the outbreak. Phylogenetic analysis has revealed that HCoV-19 is a novel beta-CoV distinct from SARS-CoV and MERS-CoV^1,2,4^. To date, the most closely related virus to HCoV-2019 is RaTG13, identified from a *Rhinolophus affinis* bat sampled in Yunnan province in 2013^2^. This virus shared 96.1% nucleotide identity and 92.9% identity in the S gene, again suggesting that bats play a key role as coronavirus reservoirs^2^. Notably, however, two research groups recently reported several novel beta-CoVs related to HCoV-19 in Malayan pangolins (*Manis javanica*) that were illegally imported into Guangxi (GX) and Guangdong (GD) provinces, southern China^7,8^. Although these pangolins CoVs are more distant to HCoV-19 than RaTG13 across the virus genome as a whole, they are very similar to HCoV-19 in the receptor binding domain (RBD) of the S protein, including at the amino acid residues thought to mediate binding to ACE2^8^. It is therefore possible that pangolins play an important role in the ecology and evolution of CoVs, although whether they act as intermediate hosts for HCoV-19 is currently unclear. Indeed, the discovery of viruses in pangolins suggests that there is a wide diversity of CoVs still to be sampled in wildlife, some of which may be directly involved in the emergence of HCoV-19.

Between May and October, 2019, we collected a total of 302 samples from 227 bats from Mengla County, Yunnan Province in southern China (Extended Data Table 1). These bats belonged to 20 different species, with the majority of samples from *Rhinolophus malayanus* (n=48, 21.1%), *Hipposideros larvatus* (n=41, 18.1%) and *Rhinolophus stheno* (n=39, 17.2%). The samples comprised multiple tissues, including patagium (n=219), lung (n=2) and liver (n=3), and feces (n=78). All but three bats were sampled alive and subsequently released. Based on the bat species primarily identified according to morphological criteria and confirmed through DNA barcoding, the 224 tissues and 78 feces were merged into 38 and 18 pools, respectively, with each pool including 1 to 11 samples of the same type (Extended Data Table 1). These pooled samples were then used for next generation sequencing (NGS).

Using next-generation metagenomic sequencing we successfully obtained 11954 and 64224 reads in pool No. 39 (from a total of 78,477,464 clean reads) that mapped to a SARS-like bat coronavirus, Cp/Yunnan2011^9^ (JX993988), and to HCoV-19. From this, we generated two preliminary consensus sequences. Pool 39 comprised 11 feces from *Rhinolophus malayanus* collected between May 6 and July 30, 2019. After a series of verification steps, including re-mapping and Sanger sequencing (Extended Data Table 2 and Figures 1-3), one partial (23395 bp) and one complete (29671 bp) beta-CoV genome sequences were obtained and termed BetaCoV/Rm/Yunnan/YN01/2019 (RmYN01) and BetaCoV/Rm/Yunnan/YN02/2019 (RmYN02), respectively. Notably, 20 positions in the RmYN02 genome displayed nucleotide polymorphisms in the NGS data, although these did not include the S1/S2 cleavage site (Extended Data Figure 3). Only a few reads in the remaining 55 pools could be mapped to the reference CoV genomes. The sequence identity between RmYN01 and Cp/Yunnan2011 across the aligned regions was 96.9%, whereas that between RmYN01 and HCoV-19 was only 79.7% across the aligned regions and 70.4% in the spike gene.

In contrast, RmYN02 was closely related to HCoV-19, exhibiting 93.3% nucleotide sequence identity, although it was less similar to HCoV-19 than RaTG13 (96.1%) across the genome as a whole (Fig. 1a). RmYN02 and HCoV-19 were extremely similar (>96% sequence identity) in most genomic regions (e.g. 1ab, 3a, E, 6, 7a, N and 10) (Fig. 1a). In particular, RmYN02 was 97.2% identical to HCoV-19 in the longest encoding gene region, 1ab (n=21285). However, RmYN02 exhibited far lower sequence identity to HCoV-19 in the S gene (nucleotide 71.8%, amino acid 72.9%), compared to 97.4% amino acid identity between RaTG13 and HCoV-19 (Fig. 1a). Strikingly, RmYN02 only possessed 62.4% amino acid identity to HCoV-19 in the RBD, whereas the pangolin beta-CoV from Guangdong had amino acid identity of 97.4%^7^, and was the closest relative of HCoV-19 in this region. A similarity plot estimated using Simplot^10^ also revealed that RmYN02 was more similar to HCoV-19 than RaTG13 in most genome regions (Fig. 1b). Again, in the RBD, the pangolin/MP789/2019 virus shared the highest sequence identity to HCoV-19 (Fig. 1c).

**Fig. 1.**
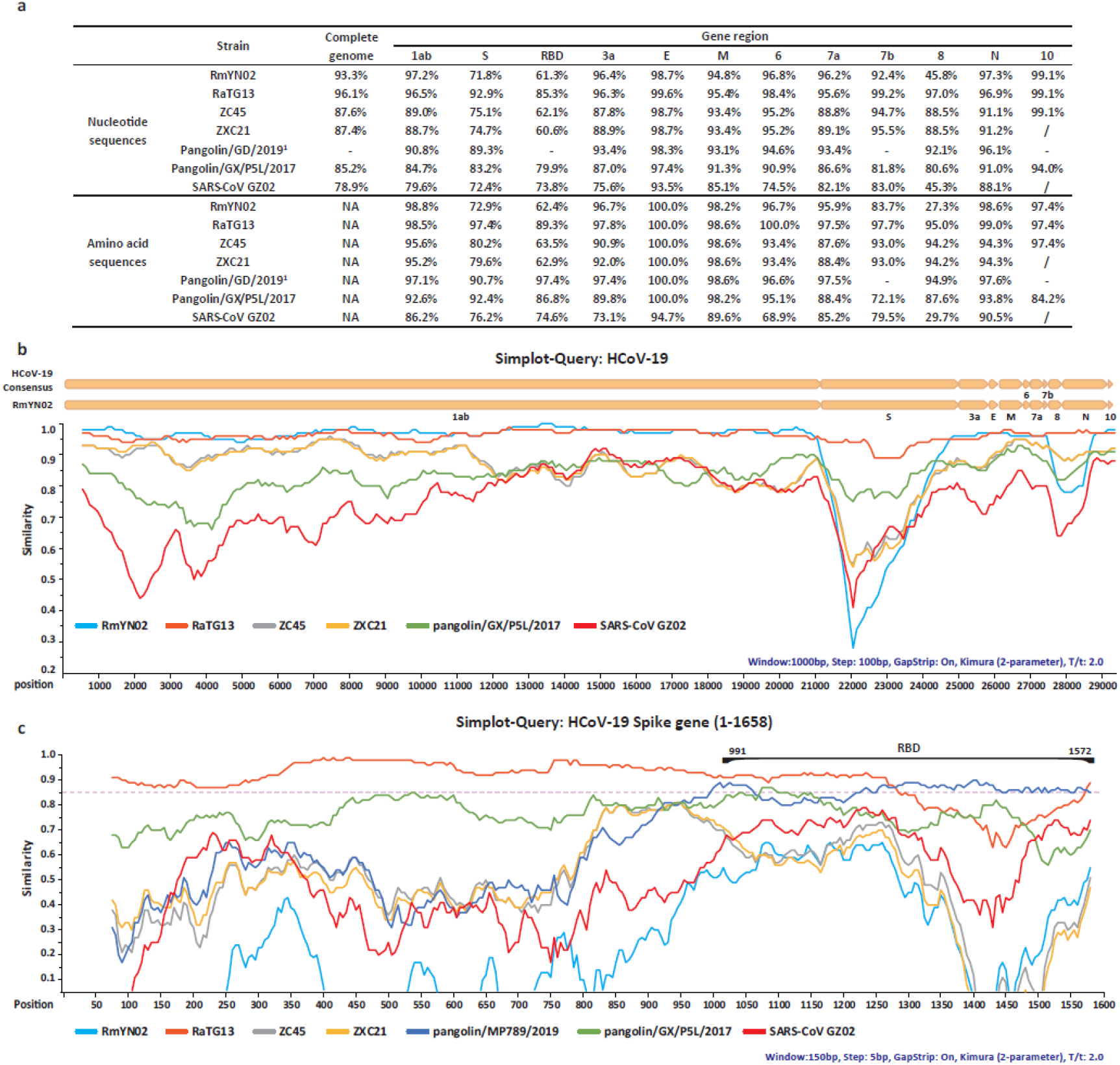
Patterns of sequence identity between the consensus sequences of HCoV-19 and representative beta-CoVs. (a) Sequence identities for HCoV-19 compared to representative beta-CoVs, including RmYN02, RaTG13 (EPI_ISL_402131), ZC45 (MG772933), ZXC21 (MG772934), pangolin/GX/P5L/2017 (EPI_ISL_410540) and SARS-CoV GZ02 (AY390556). ^1^Pangolin/GD/2019 represents a merger of GD/P1L and GD/P2S, and these values were adapted from the reference^7^. “-”: No corresponding values in reference^7^. “/”: This orf is not found. (b) Whole genome similarity plot between HCoV-19 and representative viruses listed in panel (a). The analysis was performed using Simplot, with a window size of 1000bp and a step size of 100bp. (c) Similarity plot in the spike gene (positions 1-1658) between HCoV-19 and representative viruses listed in panel (a). The analysis was performed using Simplot, with a window size of 150bp and a step size of 5bp.

Results from both homology modelling^1^, *in vitro* assays^2^ and resolved three-dimensional structure of the S protein^11^ have revealed that like SARS-CoV, HCoV-19 could also use ACE2 as a cell receptor. We analyzed the RBD of RmYN02, RaTG13, and the two pangolin beta-CoVs using homology modelling (Fig. 2a-2f and Extended Data Figure 4 for sequence alignment). The amino acid deletions in RmYN02 RBD made two loops near the receptor binding site that are shorter than those in HCoV-19 RBD (Fig. 2a and 2f). Importantly, the conserved disulfide bond in the external subdomain of SARS-CoV (PDB: 2DD8)^12^, HCoV-19 (PDB: 6LZG), RaTG13 (Fig. 2b), pangolin/MP789/2019 (Fig. 2c) and pangolin/GX/P5L/2017 (Fig. 2d) was missing in RmYN02 (Fig. 2f). We speculate that these deletions may cause conformational variations and consequently reduce the binding of RmYN02 RBD with ACE2 or even cause non-binding. It is possible that the bat SARS-related CoVs with loop deletions, including RmYN02, ZXC21 and ZC45, use a currently unknown receptor. In contrast, RaTG13 (Fig. 2b), pangolin/ MP789/2019 (Fig. 2c) and pangolin/P5L/2017 (Fig. 2d) did not have the deletions, and had similar conformations at their external domains, indicating that they may also use ACE2 as cell receptor although, with the exception of pangolin/MP789/2019 (see below), all exhibited amino acid variation to HCoV-19. Indeed, the pangolin/MP789/2019 virus showed highly structural homology with HCoV-19 (Fig. 2e).

**Fig. 2.**
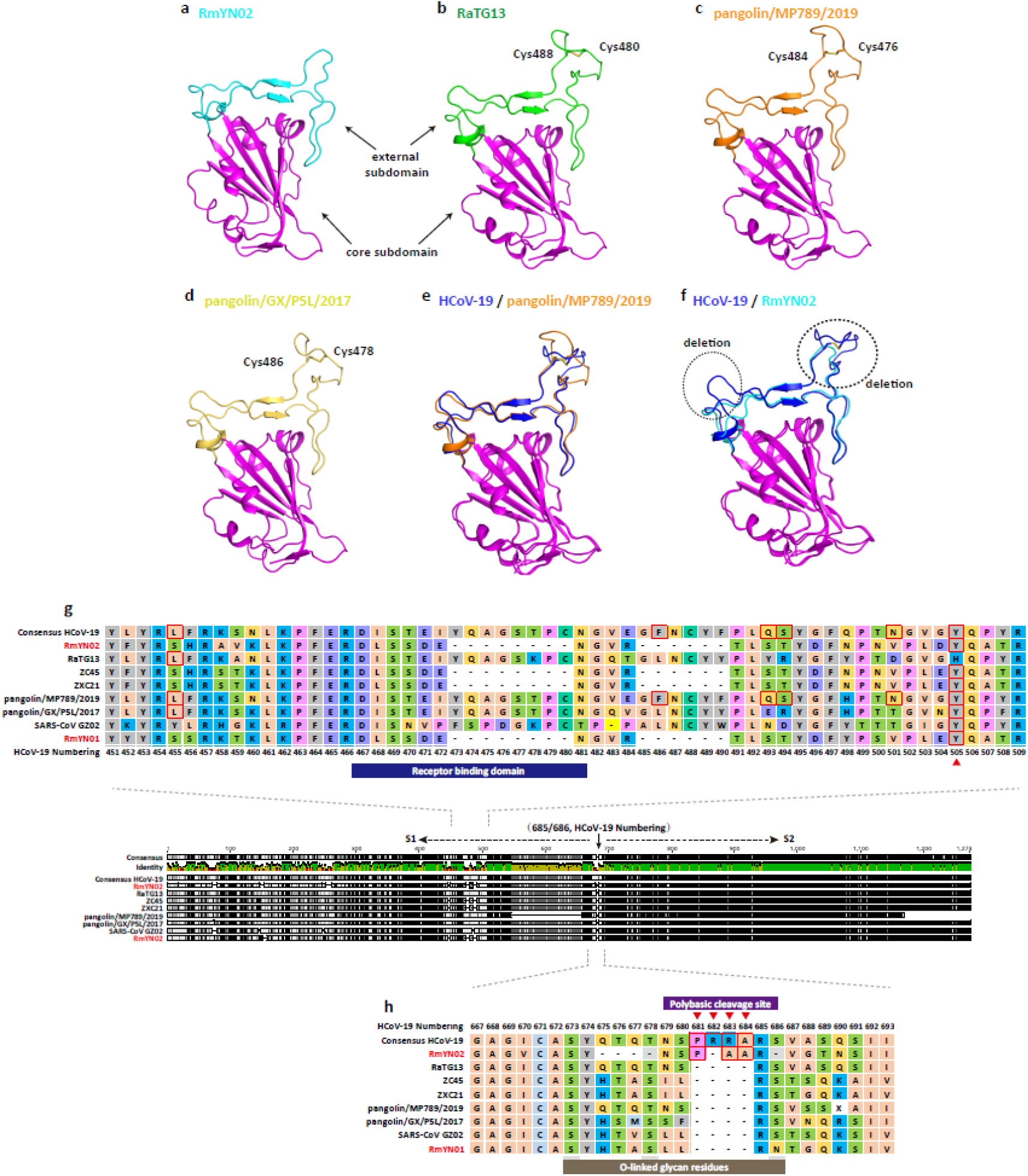
Homology modelling of the RBD structures and molecular characterizations of the S1/S2 cleavage site of RmYN02 and representative beta-CoVs. (a-d) Homology modelling and structural comparison of the RBD structures of RmYN02 and representative beta-CoVs, including (a) RmYN02, (b) RaTG13, (c) pangolin/MP789/2019 and (d) pangolin/GX/P5L/2017. The three-dimensional structures of the RBD from Bat-SL-CoV RmYN02, RaTG13, pangolin/MP789/2019 and pangolin/GX/P5L/2017 were modeled using the Swiss-Model program^21^ employing the RBD of SARS-CoV (PDB: 2DD8) as a template. All the core subdomains are colored magenta, and the external subdomains of RmYN02, RaTG13, pangolin/MP789/2019 and pangolin/GX/P5L/2017 are colored cyan, green, orange and yellow, respectively. The conserved disulfide bond in RaTG13, pangolin/GD and pangolin/GX is highlighted, while it is missing in RmYN02 due to a sequence deletion. (e-f) Superimposition of the RBD structure of pangolin/MP789/2019 (e) and RmYN02 (f) with that of HCoV-19. The two deletions located in respective loops in RmYN02 are highlighted using dotted cycles. (g) Molecular characterizations of the RBD of RmYN02 and the representative beta-CoVs. (h) Molecular characterizations of the cleavage site of RmYN02 and the representative beta-CoVs.

Six amino acid residues at the RBD (L455, F486, Q493, S494, N501 and Y505) have been reported to be major determinants of efficient receptor binding of HCoV-19 to ACE2^13^. As noted above, and consistent with the homology modelling, pangolin/MP789/2019 possessed the identical amino acid residues to HCoV-19 at all six positions^7^. In contrast, both RaTG13, RmYN02 and RmYN01 possessed the same amino acid residue as HCoV-19 at only one of the six positions each (RaTG13, L455; RmYN02, Y505; RmYN01, Y505) (Fig. 2g), despite RaTG13 being the closest relative in the spike protein. Such an evolutionary pattern is indicative of a complex combination of recombination and natural selection^7,14^.

The S protein of CoVs is functionally cleaved into two subunits, S1 and S2^15^ in a similar manner to the haemagglutinin (HA) protein of avian influenza viruses (AIVs). The insertion of polybasic amino acids at the cleavage site in the HAs of some AIV subtypes is associated with enhanced pathogenicity^16,17^. Notably, HCoV-19 is characterized by a four-amino-acid-insertion at the junction of S1 and S2, not observed in other lineage B beta-CoVs^18,19^. This insertion, which represents a poly-basic (furin) cleavage site, is unique to HCoV-19 and is present in all HCoV-19 sequenced so far. The insertion of three residues, PAA, at the junction of S1 and S2 in RmYN02 (Fig. 2h and Extended Data Figure 2) is therefore of major importance. Although the inserted residues (and hence nucleotides) are not the same as those in RmYN02, and hence are indicative of an independent insertion event, that they are presented in wildlife (bats) strongly suggests that they are of natural origin and have likely acquired by recombination. As such, these data are strongly suggestive of a natural zoonotic origin of HCoV-19.

We next performed a phylogenetic analysis of RmYN02, RaTG13, HCoV-19 and the pangolin beta-CoVs. Consistent with a previous research^7^, the pangolin beta-CoVs formed two well-supported sub-lineages, representing animal seized by anti-smuggling authorities in Guangxi (Pangolin-CoV/GX) and Guangdong (Pangolin-CoV/GD) provinces (Fig. 3a and Extended Data Figure 5). However, whether pangolins are natural reservoirs for these viruses, or they acquired these viruses independently from bats or other wildlife, requires further sampling^7^. More notable was that RmYN02 was the closest relative of HCoV-19 in most of the virus genome, although these two viruses were still separated from each other by a relatively long branch length (Fig. 3a and Extended Data Figure 5). In the spike gene tree, HCoV-19 clustered with RaTG13 and was distant from RmYN02, suggesting that the latter virus has experienced recombination in this gene (Fig. 3b and Extended Data Figure 6). In phylogeny of the RBD, HCoV-19 was most closely related to pangolin-CoV/GD, with the bat viruses falling in more divergent positions, again indicative of recombination (Fig. 3c and Extended Data Figure 7). Finally, phylogenetic analysis of the complete RNA dependent RNA polymerase (RdRp) gene, which is often used in the phylogenetic analysis of RNA viruses, revealed that RmYN02, RaTG13 and HCoV-19 formed a well-supported sub-cluster distinct from the pangolin viruses (Fig. 3d and Extended Data Figure 8).

**Fig. 3.**
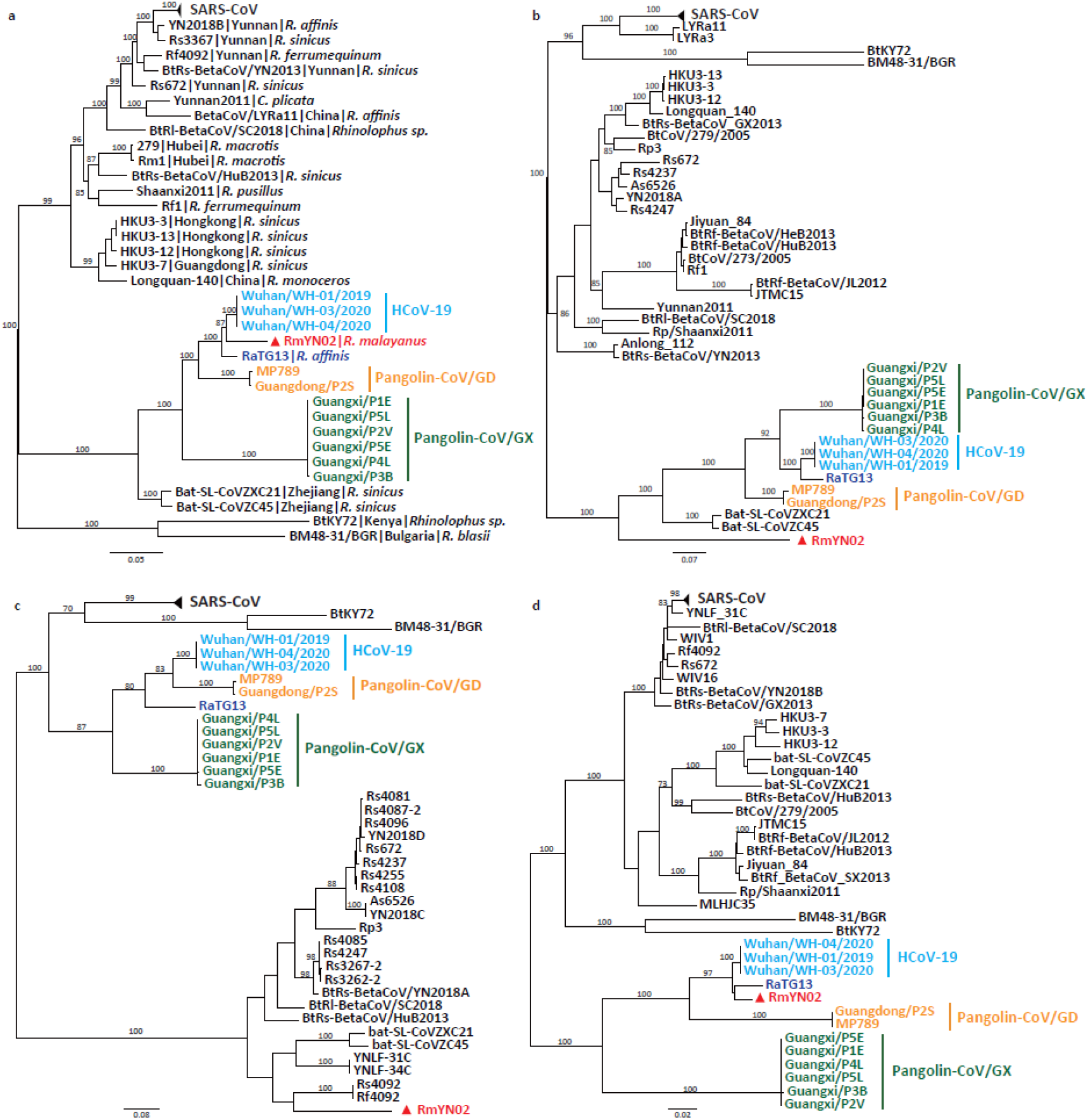
Phylogenetic analysis of HCoV-19 and representative viruses from the subgenus *Sarbecoronavirus*. (a) Phylogenetic tree of the full-length virus genome. (b) the S gene. (c) the RBD. (d) the RdRp. Phylogenetic analysis was performed using RAxML^22^ with 1000 bootstrap replicates, employing the GTR nucleotide substitution model. RBD is delimited as the gene region 991-1572 of the spike gene according to the reference^7^. All the trees are midpoint rooted for clarity.

We confirmed the bat host of RmYN02, *Rhinolophus malayanus*, by analyzing the sequence of the cytochrome b (*Cytb*) gene from the next generation sequencing data; this revealed 100% sequence identity to a *Rhinolophus malayanus* isolate (GenBank accession MK900703). Both *Rhinolophus malayanus* and *Rhinolophus affinis* are widely distributed in southwest China and southeast Asia. Generally, they do not migrate over long distances and are highly gregarious such that they are likely to live in the same caves, which might facilitate the exchange of viruses between them and the occurrence of recombination. Notably, RaTG13 was identified from anal swabs and RmYN02 was identified from feces, which is a simple, but feasible way for bats to spread the virus to other animals, especially species that can utilize cave environments.

Based on the currently available data we propose that HCoV-19 likely originates from multiple naturally occurring recombination events in wildlife. A virus from bats likely provides the genetic backbone of HCoV-19, with further recombination events with bats and perhaps other wildlife species resulting in the acquisition of the Spike protein, RBD and the polybasic cleavage site. Similar recombination events have been also implicated in the origin of SARS-CoV^20^, although it is clear that a far wider sampling of wildlife will be required to reveal the exact species involved and the exact series of recombination events.

