## Supplementary materials for "A novel bat coronavirus reveals natural insertions at the S1/S2 cleavage site of the Spike protein and a possible recombinant origin of HCoV-19"

#### Methods

##### Sample collection

Between May and October, 2019, a total of 302 samples from 227 bats were collected from Mengla County, Yunnan Province in southern China (Extended Data Table 1). These bats belonged to 20 different species, with the majority of samples belonging to *Rhinolophus malayanus* (n=48, 21.1%), *Hipposideros larvatus* (n=41, 18.1%) and *Rhinolophus stheno* (n=39, 17.2%). The samples included patagium (n=219), lung (n=2) and liver (n=3), and feces (n=78), all but three bats were sampled alive and subsequently released. All samples were first stored in RNAlater and then kept at -80°C until use.

##### Next generation sequencing

Based on the bat species primarily identified according to morphological criteria and confirmed through DNA barcoding, the 224 tissue and 78 fecal samples were merged into 38 and 18 pools, respectively, with each pool including 1 to 11 samples of the same type (Extended Data Table 1). Samples were transferred into the RNAiso Plus reagent (TAKARA) for homogenization with steel beads. Total RNA was extracted and subsequently purified using EZNA Total RNA Kit (OMEGA). Libraries were constructed using the NEB Next Ultra RNA Library Prep Kit (NEB). rRNA of feces or tissues was removed using the TransNGS rRNA Depletion (Bacteria) Kit and TransNGS rRNA Depletion (Human/Mouse/Rat) Kit (TransGen), respectively. Paired-end (150 bp) sequencing of each RNA library was performed on the NovaSeq 6000 platform (Illumina) carried out by Novogene Bioinformatics Technology (Beijing, China).

#### Genome assembly and annotation

Raw reads were obtained from the 56 pools and were then adaptor- and quality- trimmed with the Fastp program[1]. The clean reads were then mapped to reference genomes of representative CoV genomes using Bowtie 2[2], including HCoV-19 (MDC60013002-01), SARS-CoV (AY508724, AY485277, AY390556 and AY278489), SARS-like-CoV (DQ084200, DQ648857, GQ153542, GQ153547, JX993987, JX993988, KF294455, KF294457, KJ473814, KJ473815, KJ473816, KT444582, KY417142, KY417145, KY417146, KY417148, KY417151, KY417152, KY770859, MK211374, MK211376, and MK211377), ZC45 (MG772933), ZXC21 (MG772934), MERS-CoV (JX869059), other representative beta-CoV genomes (AY391777, EF065505, EF065509, EF065513, FJ647223, KC545386, KF636752, KM349744, KU762338, and MK167038) and alphacoronavirus genomes (NC_002645, AY567487). 11954 and 64224 reads in pool No. 39 (a total of 78,477,464 clean reads) were mapped to both a bat coronavirus Cp/Yunnan2011 (JX993988)[3] and HCoV-19, generating two preliminary consensus sequences, termed BetaCoV/Rm/Yunnan/YN01/2019 (RmYN01) and BetaCoV/Rm/Yunnan/YN02/2019 (RmYN02), respectively. However, there were only few reads in the remaining 55 pools that could be mapped to these reference CoV genomes. Pool 39 comprised 11 feces from *Rhinolophus malayanus* collected between May 6 and July 30, 2019.

To validate the two novel CoV genomes, the clean reads of pool 39 were then *de novo* assembled using Trinity[4] with default settings. The assembled contigs were compared with the consensuses obtained in the previous step and merged using Geneious (version 11.1.5) (https://www.geneious.com). We found that contigs with high and low abundance corresponded to RmYN02 and RmYN01, respectively, with the abundance of RmYN02 5-10 times higher than that of RmYN01. The gaps between contigs of RmYN02 were complemented by re-mapping the reads to the ends of the contigs, which produced the full-length genome sequence of RmYN02. However, due to the limited number of reads available, only a partial genome sequence of RmYN01 was obtained (23395 bp). Reads were then mapped to the full-length genome sequence of RmYN02 using Bowtie 2 to check base consistency at each nucleotide site. Moreover, to perform *de novo* assembly for pool 39 and to further distinguish RmYN01 and RmYN02, PeHaplo[5] was used with various overlap parameters (70, 80, 100, and 120). The consensus of each run of PeHaplo was then compared, generating the final full-length genome of RmYN02 (29671 bp). The sequence identity between RmYN01 and Cp/Yunnan2011 across the aligned regions was 96.9%, whereas that between RmYN01 and HCoV-19 was only 79.7%.

#### Bioinformatics analyses

Reference virus genomes were obtained from NCBI/GenBank (https://www.ncbi.nlm.nih.gov/) using Blastn with HCoV-19 as a query. The beta-CoVs from pangolins (Extended Data Table 3) were retrieved from GISAID (www.gisaid.org/). The open reading frames (ORFs) of the verified genome sequences were predicted using Geneious (version 11.1.5). Pairwise sequence identities were also calculated using Geneious. Potential recombination events were investigated using Simplot (version 3.5.1)[6].

The three-dimensional structures of RBD from RmYN02, RaTG13, pangolin/GD and pangolin/GX were modeled using Swiss-Model program[7] using SARS CoV RBD structure (PDB: 2DD8)[8] as a template.

Multiple sequence alignment of HCoV-19 and the reference sequences was performed using Mafft[9]. Phylogenetic analyses of the complete genome and major encoding regions were performed using RAxML[10] with 1000 bootstrap replicates, employing the GTR nucleotide substitution model (Fig. 3). Phylogenetic analysis was also performed using MrBayes[11], employing the GTR nucleotide substitution model (Extended Data Figures 5-8). Ten million steps were run, with trees and parameters sampled every 1,000 steps.

#### Sanger sequencing

Based on the spike gene sequence of RmYN02, a TaqMan-based qPCR was performed to test the feces of pool 39 (Extended Data Table 2). Pool 39 comprised 11 feces from *Rhinolophus malayanus* collected between May 6 and July 30, 2019. However, only eight original samples were left after NGS. The results indicated that the fecal sample No. 123 from *R. malayanus*, collected on June 25$^{th}$, 2019, was positive for RmYN02 (Extended Data Figure 1). To further confirm the S1/S2 cleavage site and the 1b (RdRp) gene sequence of RmYN02, five pair primers, F1/R1-F4/R4 and F6/R6, were designed for Sanger sequencing (Extended Data Table

2). The consensus gene sequence of Sanger sequencing of the amplified products was consistent with those from NGS (Extended Data Figure 2).

### Data availability

The sequences of RmYN01 and RmYN02 have also deposited in the GISAID with accession numbers: EPI_ISL_412976 and EPI_ISL_412977.

### Acknowledgments

This work was supported by the Academic Promotion Programme of Shandong First Medical University (2019QL006 and 2019PT008), the Strategic Priority Research Programme of the Chinese Academy of Sciences (XDA19090118, XDB29010102 and XDA20050202), the Chinese National Natural Science Foundation (U1602265), the National Major Project for Control and Prevention of Infectious Disease in China (2017ZX10104001-006), and the High-End Foreign Experts Program of Yunnan Province (Y9YN021B01). W.S. was supported by the Taishan

Scholars Programme of Shandong Province (ts201511056). Y.B. is supported by the NSFC Outstanding Young Scholars (31822055) and Youth Innovation Promotion Association of CAS (2017122). E.C.H. is supported by an ARC Australian Laureate Fellowship (FL170100022). We thank all the scientists, especially Professor Wuchun Cao and Professor Yi Guan, who kindly shared their genomic sequences of the coronaviruses used in this study.

### Author contributions

W.S., Y.B. and A.C.H. designed and supervised research. X.C. and A.C.H. collected the samples. H.Z. and Y.L. processed the samples. H.T. and J.L. performed genome assembly and annotation. H.Z., J.L. and T.H. performed the genome analysis and interpretation. W.S., Y.B. and A.C.H. wrote the paper. X.C., P.W., D.L., J.Y. and E.C.H. assisted in data interpretation and edited the paper.

### Competing interests

The authors declare no competing interests.

### Extended Data

**Extended Data Figure 1. The detection result of the eight fecal samples of pool 39 for RmYN02 using real-time PCR primers and Taqman probe.** Two replicates were set for each original sample.

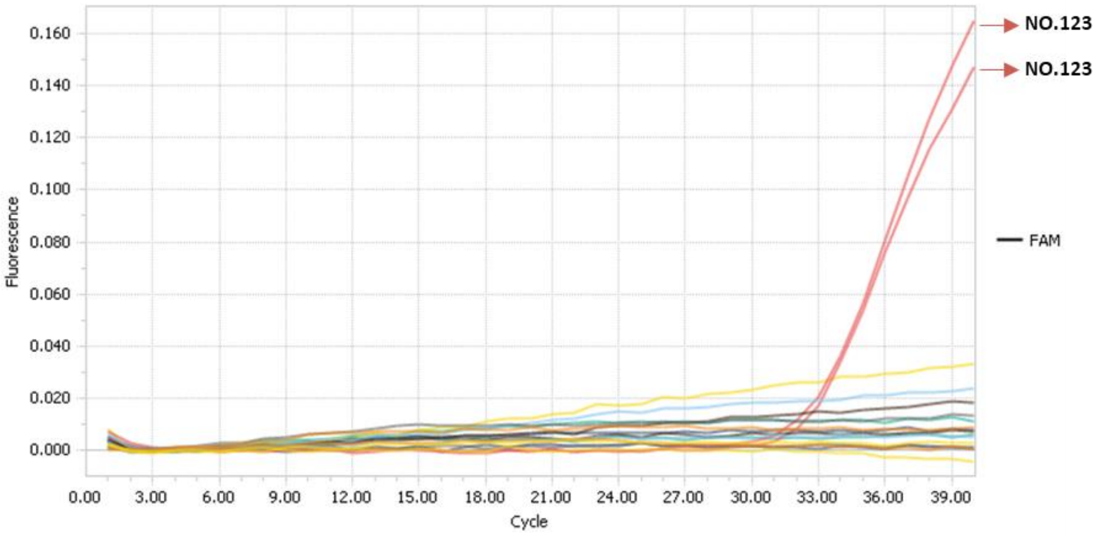

**Extended Data Figure 2. Comparison of NGS consensus and Sanger sequencing of the RBD and the cleavage site of RmYN02.** The insertion of the multiple amino acids at the S1/S2 cleavage site is highlighted.

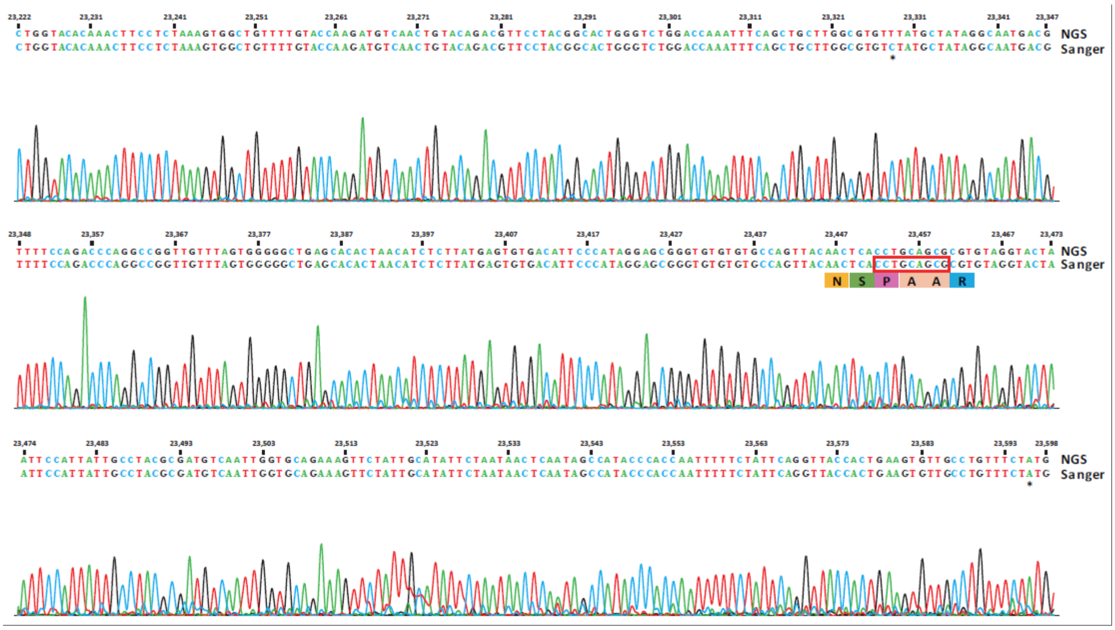

**Extended Data Figure 3. Sites in the RmYN02 genome that display nucleotide polymorphisms in the NGS data.** The positions of these sites in RmYN02 were provided at the bottom of the figure. “/”: Sanger sequencing was not performed.

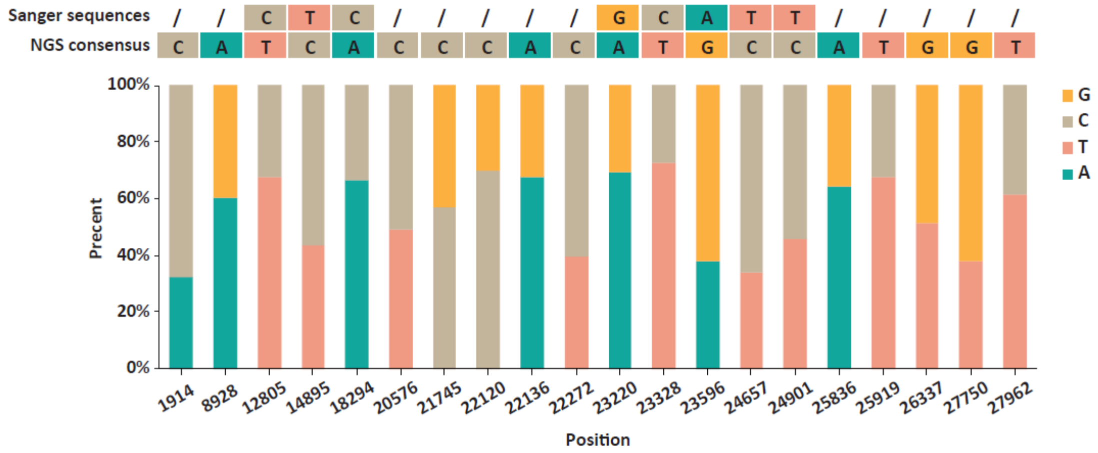

**Extended Data Figure 4. Sequence alignment of the RBDs from RmYN02 and representative beta-CoVs.** The blue triangles indicate amino acids from the SARS-CoV S protein that impact binding to ACE2. The yellow rectangles highlight the three deletions in sequence resulting two loops shorten in RmYN02.

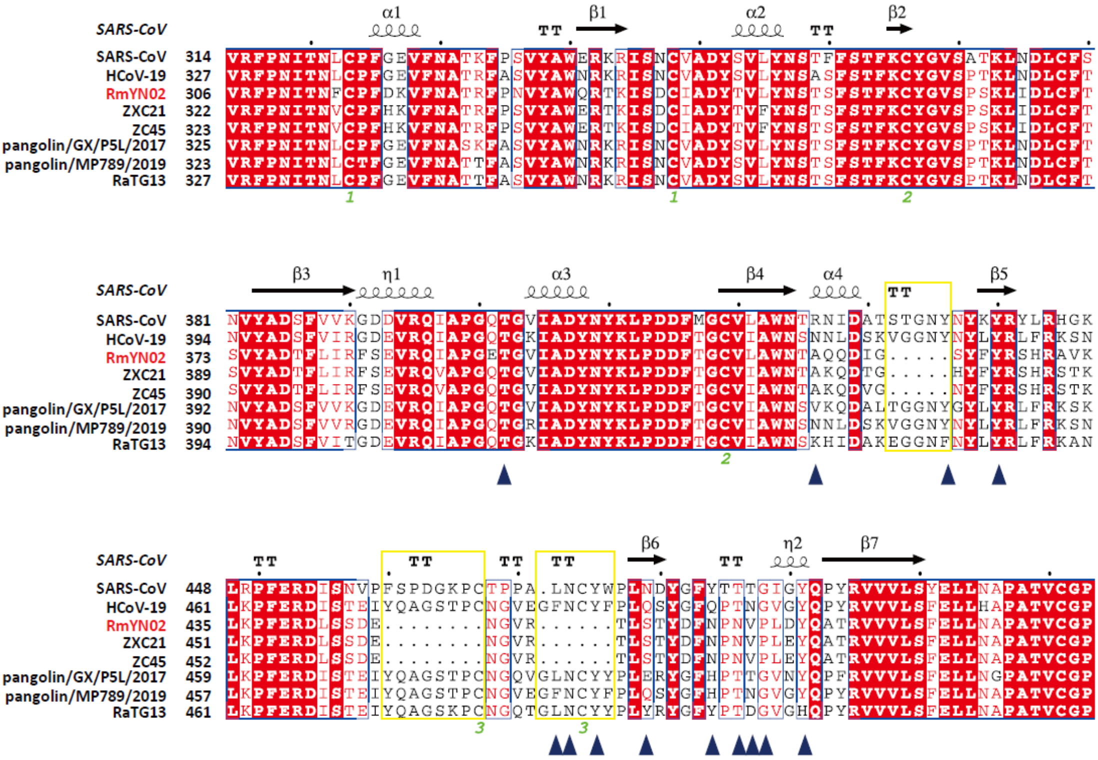

**Extended Data Figure 5. Bayesian phylogenetic analysis of the full-length virus genome of HCoV-19 and representative viruses of the subgenus *Sarbecoronavirus*.** Phylogenetic analysis was also performed using MrBayes, employing the GTR nucleotide substitution model. Ten million steps were run, with trees and parameters sampled every 1,000 steps. The tree is midpoint rooted for clarity.

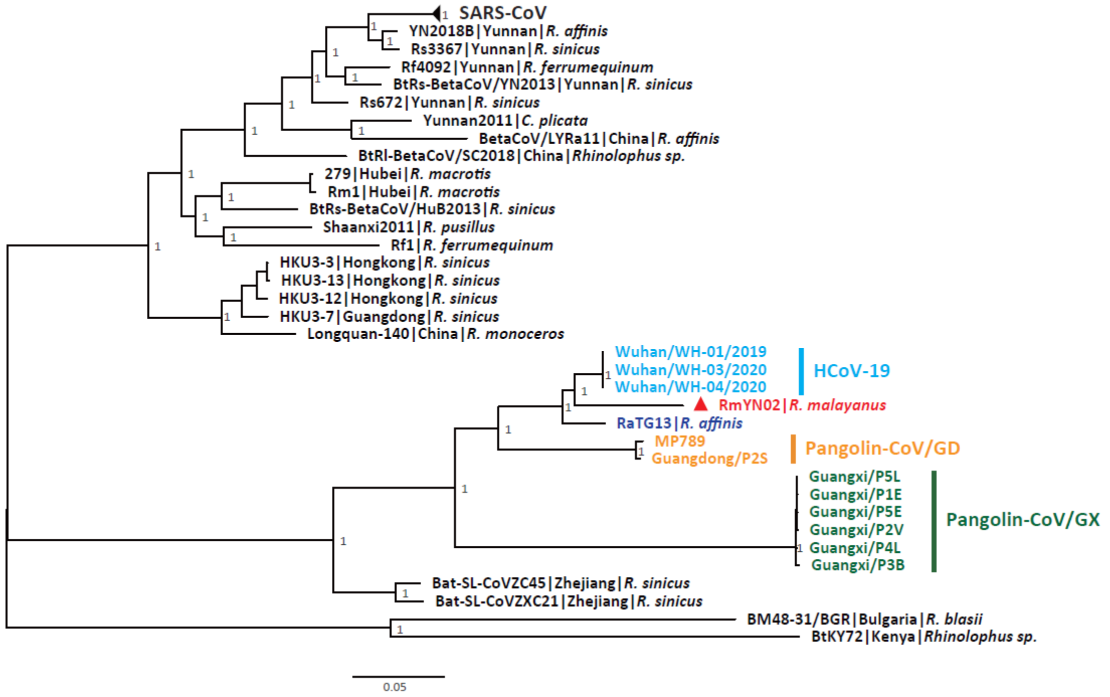

**Extended Data Figure 6. Bayesian phylogenetic analysis of the spike gene of HCoV-19 and representative viruses of the subgenus *Sarbecoronavirus*.** Phylogenetic analysis was also performed using MrBayes, employing the GTR nucleotide substitution model. Ten million steps were run, with trees and parameters sampled every 1,000 steps. The tree is midpoint rooted for clarity.

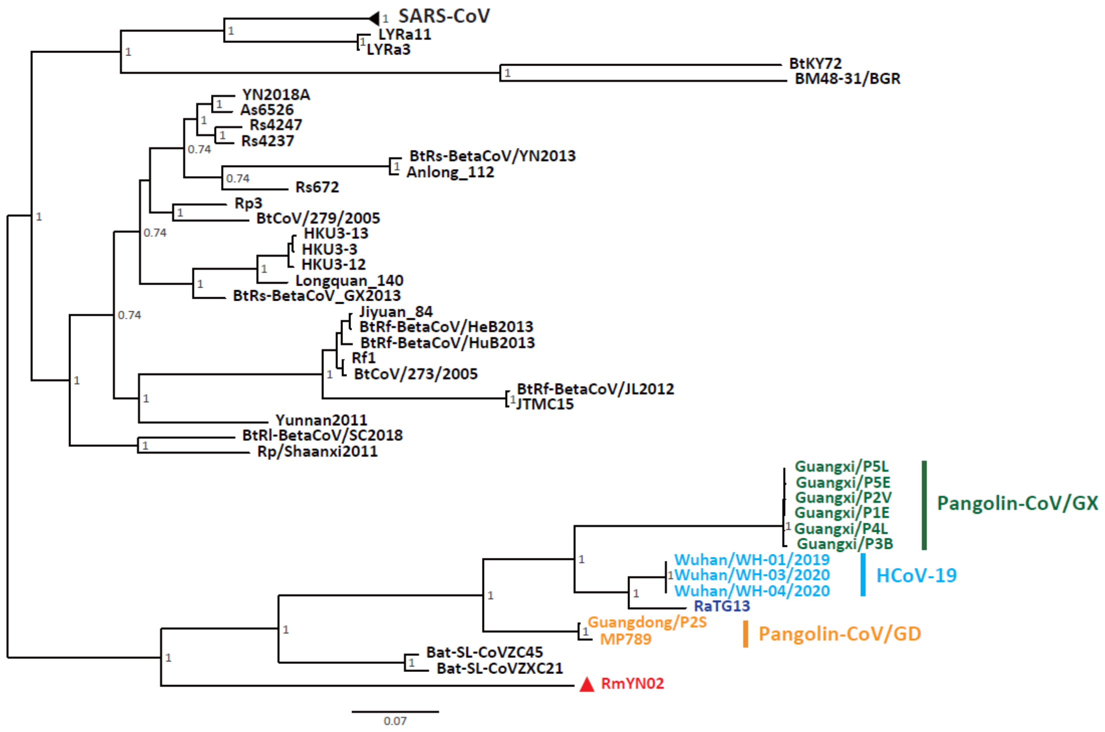

**Extended Data Figure 7. Bayesian phylogenetic analysis of the RBD of HCoV-19 and representative viruses of the subgenus *Sarbecoronavirus*.** RBD is delimited as the gene region 991-1572 of the spike gene according to the reference[7]. Phylogenetic analysis was also performed using MrBayes, employing the GTR nucleotide substitution model. Ten million steps were run, with trees and parameters sampled every 1,000 steps. The tree is midpoint rooted for clarity.

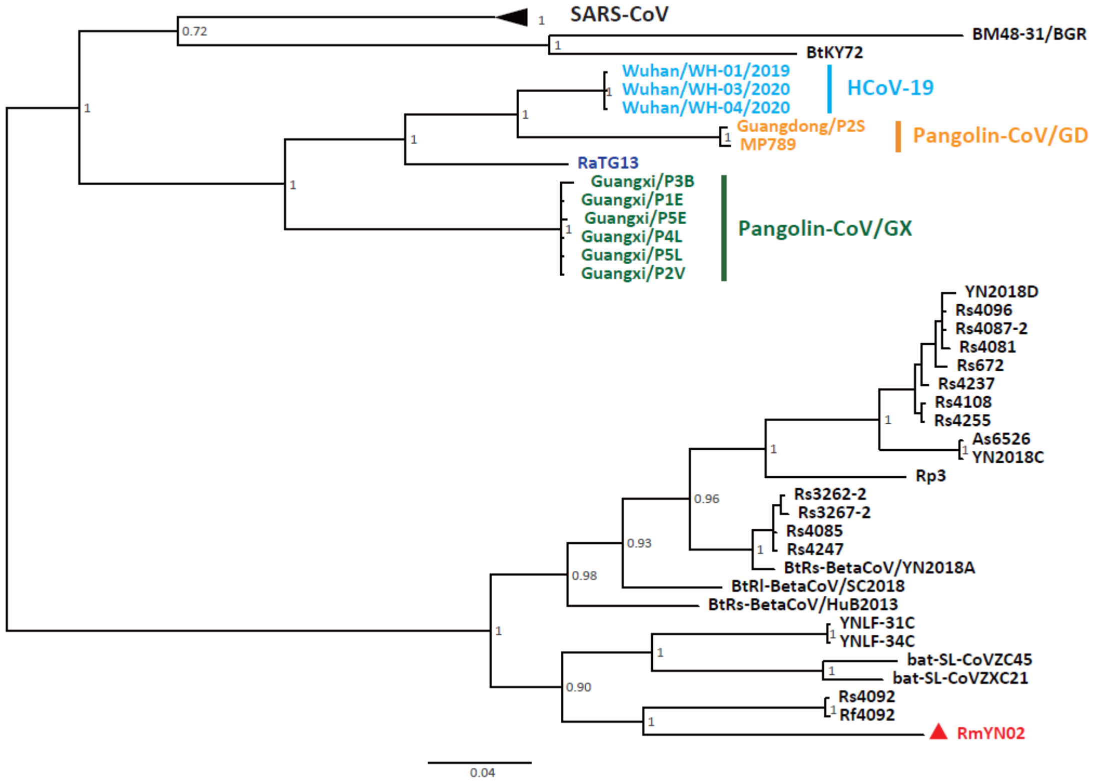

**Extended Data Figure 8. Bayesian phylogenetic analysis of the RdRp gene of HCoV-19 and representative viruses of the subgenus *Sarbecoronavirus*.** Phylogenetic analysis was also performed using MrBayes, employing the GTR nucleotide substitution model. Ten million steps were run, with trees and parameters sampled every 1,000 steps. The tree is midpoint rooted for clarity.

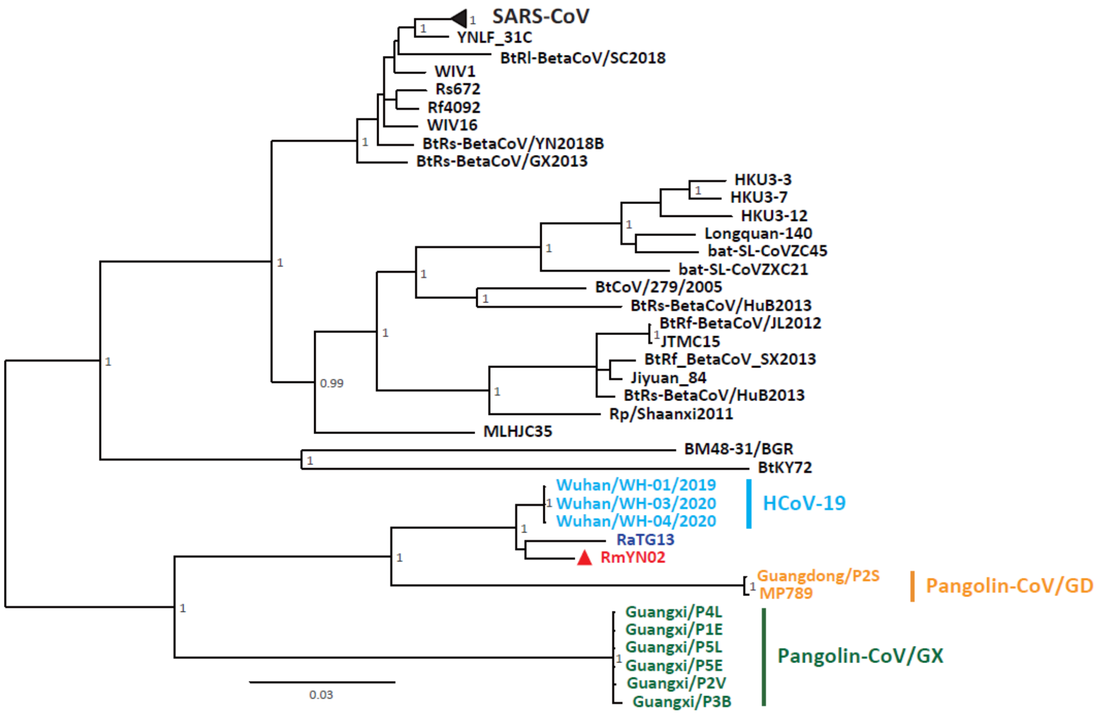

**Extended Data Table 1. Summary of the bat samples collected in the present study and the pooling strategy used for next-generation sequencing.**

| Species | Individual number | | | | | | | Number of Libraries | Sample number | | | | |
|---|---|---|---|---|---|---|---|---|---|---|---|---|---|
| | May | Jun | Jul | Aug | Sep | Oct | May-Oct | | Samples per library | Patagium | Lung | Liver | Feces |
| *Rhinolophus malayanus*[a] | 6 | 24 | 9 | | | | 39 | 5 | 10, 10, 8, 9, 11 | 37 | | | 11 |
| *Rhinolophus stheno*[a] | 14 | 12 | 16 | 6 | | | 48 | 7 | 10, 5, 10, 10, 9, 7, 9 | 44 | | | 16 |
| *Hipposideros larvatus (complex)*[a] | 4 | 8 | 12 | 5 | 7 | 5 | 41 | 8 | 10, 10, 10, 10, 7, 7, 1, 1 | 40 | 1 | 1 | 14 |
| *Rhinolophus sinicus*[a] | 4 | 4 | 4 | 3 | 2 | | 17 | 3 | 10, 7, 8 | 17 | | | 8 |
| *Myotis laniger*[a] | 7 | | 2 | | | | 9 | 1 | 9, | 9 | | | |
| *Rhinolophus siamensis*[a] | 1 | 3 | 1 | 2 | | 1 | 8 | 2 | 8, 3 | 8 | | | 3 |
| *Hipposideros pomona*[a] | 1 | | 4 | 7 | 1 | | 13 | 3 | 8, 5, 8 | 13 | | | 7 |
| *Kerivoula hardwickii*[a] | 1 | | | 1 | 1 | | 3 | 1 | 3, | 3 | | | 1 |
| *Murina cyclotis*[a] | 1 | 2 | 2 | 2 | | | 7 | 2 | 7, 3 | 7 | | | 3 |
| *Aselliscus stoliczkanus*[a] | 1 | | | 2 | 1 | 1 | 5 | 2 | 5, 3 | 5 | | | 3 |
| *Myotis muricola*[a] | | 3 | 1 | 2 | 1 | 1 | 8 | 1 | 8, | 8 | | | |
| *Kerivoula sp.* [a] | 2 | | | | | | 2 | 2 | 2, 1 | 2 | | | 1 |
| *Rhinolophus paradoxolophus* [a] | 1 | | | | | | 1 | 2 | 1, 1 | 1 | | | 1 |
| *Kerivoula papillosa*[a] | | | 1 | 1 | | | 2 | 1 | 2, | 2 | | | |

| | | | | | | | | | | | | | |
|---|---|---|---|---|---|---|---|---|---|---|---|---|---|
| *Tylonycteris robustula*[a] | | 1 | | | | | 1 | 1 | 1, | 1 | | | |
| *Harpiocephalus harpia*[a] | | | | 1 | | | 1 | 2 | 1, 1 | 1 | | | 1 |
| *Hipposideros armiger*[a] | | | | | 1 | 2 | 3 | 3 | 2, 1, 1 | 3 | | | 1 |
| *Rhinolophus pearsonii*[a] | | | | 3 | | 3 | 6 | 3 | 6, 2, 3 | 6 | | | 5 |
| *Chaerephon plicata*[b] | | | 4 | | | | 4 | 3 | 4, 2, 1 | 4 | | 1 | 2 |
| *Taphozous melanopogon*[b] | | | 9 | | | | 9 | 4 | 8, 1, 1, 1 | 8 | 1 | 1 | 1 |
| Total | 43 | 57 | 65 | 35 | 14 | 13 | | 56 | | 219 | 2 | 3 | 78 |

Note: [a]Samples were collected from Mengla, Xishuangbanna, Yunan (101.27156323E, 21.91889683N).

[b]Samples were collected from Mengla, Xishuangbanna, Yunan (21.5932019N, 101.2200914E).

Different colors represent different sample types. Patagium, black; Lung, purple; Liver, blue; Feces, red.

**Extended Data Table 2. Oligonucleotide primers designed to detect RmYN02 and to amplify the spike and 1b genes.**

| Primers (nucleotide positions)[a] | Sequence, 5'→3' | Product length, bp |
|---|---|---|
| Forward qF (21344-21366) | ACCCAATTCAGTTGTCTTCCTAT | 146 |
| Reverse qR (21469-21489) | TCTAACGATGAGCCTACCCTT | |
| Probe qP (21411-21438) | TGCTGTTATGTCTCTTAAGGAGGGACAA | |
| Forward F6 (23070-23092) | TGGTGTTCTAACTGATTCAGATA | 2370 |
| Reverse R6 (25418-25439) | TCTCTTTTTAAGGGTTATGATT | |
| Forward F1 (12002-12024) | CTTTCCATGCAGGGTGCTGTAGA | 1488 |
| Reverse R1 (13470-13489) | CGCACGGTGTAAGACGGGCT | |
| Forward F2 (13159-13180) | TGGTCAGGCAATAACAGTTACA | 2579 |
| Reverse R2 (14332-14354) | GCACAATGCAGAATGCATCTA | |
| Forward F3 (15625-15646) | AGAGATGTTGACACAGACTTTG | 1788 |
| Reverse R3 (17310-17412) | GTACACATAGTGCTTAGCACGTA | |
| Forward F4 (17321-17345) | CAGATATAGTTGTCTTTGATGAAAT | 1937 |
| Reverse R4 (19236-19257) | AGGCAAGTTAAGGTTAGATAGC | |

[a]Values in parentheses indicate primer positions corresponding to the genome sequence of RmYN02.

**Extended Data Table 3. Acknowledgement of sharing of HCoV-19 genome sequences from the GISAID and GenBank databases.** We gratefully thank the authors listed below for sharing their genomic sequences of coronaviruses analyzed in this study.

| **Accession ID** | **Virus name** | **Location** | **Collection date** | **Originating lab** | **Submitting lab** | **Authors** |
|---|---|---|---|---|---|---|
| EPI_ISL_402131 | BetaCoV/bat/Yunnan/RaTG13/2013 | Asia / China / Yunnan / Pu'er | 2013-07-24 | Wuhan Institute of Virology, Chinese Academy of Sciences | Wuhan Institute of Virology, Chinese Academy of Sciences | Yan Zhu, Ping Yu, Bei Li, Ben Hu, Hao-Rui Si, Xing-Lou Yang, Peng Zhou, Zheng-Li Shi |
| EPI_ISL_410539 | BetaCoV/pangolin/Guangxi/P1E/2017 | Asia / China / Guangxi | 2017 | Beijing Institute of Microbiology and Epidemiology | Beijing Institute of Microbiology and Epidemiology | Wu-Chun Cao; Tommy Tsan-Yuk Lam; Na Jia; Ya-Wei Zhang; Jia-Fu Jiang; Bao-Gui Jiang |
| EPI_ISL_410541 | BetaCoV/pangolin/Guangxi/P5E/2017 | Asia / China / Guangxi | 2017 | Beijing Institute of Microbiology and Epidemiology | Beijing Institute of Microbiology and Epidemiology | Wu-Chun Cao; Tommy Tsan-Yuk Lam; Na Jia; Ya-Wei Zhang; Jia-Fu Jiang; Bao-Gui Jiang |
| EPI_ISL_410540 | BetaCoV/pangolin/Guangxi/P5L/2017 | Asia / China / Guangxi | 2017 | Beijing Institute of Microbiology and Epidemiology | Beijing Institute of Microbiology and Epidemiology | Wu-Chun Cao; Tommy Tsan-Yuk Lam; Na Jia; Ya-Wei Zhang; Jia-Fu Jiang; Bao-Gui Jiang |
| EPI_ISL_410538 | BetaCoV/pangolin/Guangxi/P4L/2017 | Asia / China / Guangxi | 2017 | Beijing Institute of Microbiology and Epidemiology | Beijing Institute of Microbiology and Epidemiology | Wu-Chun Cao; Tommy Tsan-Yuk Lam; Na Jia; Ya-Wei |

| | | | | | | |
|---|---|---|---|---|---|---|
| | | | | | | Zhang; Jia-Fu Jiang; Bao-Gui Jiang |
| EPI_ISL_ 410543 | BetaCoV/ pangolin/ Guangxi/P3B/ 2017 | Asia / China / Guangxi | 2017 | Beijing Institute of Microbiology and Epidemiology | Beijing Institute of Microbiology and Epidemiology | Wu-Chun Cao; Tommy Tsan-Yuk Lam; Na Jia; Ya-Wei Zhang; Jia-Fu Jiang; Bao-Gui Jiang |
| EPI_ISL_ 410542 | BetaCoV/ pangolin/ Guangxi/P2V/ 2017 | Asia / China / Guangxi | 2017 | Beijing Institute of Microbiology and Epidemiology | Beijing Institute of Microbiology and Epidemiology | Wu-Chun Cao; Tommy Tsan-Yuk Lam; Na Jia; Ya-Wei Zhang; Jia-Fu Jiang; Bao-Gui Jiang |
| EPI_ISL_ 410544 | BetaCoV/ pangolin/ Guangdong /P2S/2019 | Asia / China / Guangdo ng | 2019 | Beijing Institute of Microbiology and Epidemiology | Beijing Institute of Microbiology and Epidemiology | Wu-Chun Cao; Tommy Tsan-Yuk Lam; Na Jia; Ya-Wei Zhang; Jia-Fu Jiang; Bao-Gui Jiang |
| MT0840 71.1 | MP789 | China | 2019-03- 29 | | Chinese Academy of Fishery Sciences (SCSFRI, CAFS) | Jiang, J.-Z., Liu, P. and Chen, J.-P. |
